# Genetic analysis of Sephardic ancestry in the Iberian Peninsula

**DOI:** 10.1101/325779

**Authors:** Miguel Martín Álvarez-Álvarez, Neil Risch, Christopher R. Gignoux, Scott Huntsman, Elad Ziv, Laura Fejerman, Maria Esther Esteban, Magdalena Gayà-Vidal, Beatriz Sobrino, Francesca Brisighelli, Nourdin Harich, Fulvio Cruciani, Hassen Chaabani, Ángel Carracedo, Pedro Moral, Esteban González Burchard, Marc Via, Georgios Athanasiadis

**Affiliations:** Department of Evolutionary Biology, Ecology and Environmental Sciences and Biodiversity Research Institute (IRBio), Universitat de Barcelona, Barcelona, Barcelona 08028, Spain; Institute for Human Genetics, University of California San Francisco, San Francisco, CA 94118, USA; Department of Epidemiology and Biostatistics, University of California San Francisco, San Francisco, CA 94118, USA; Colorado Center for Personalized Medicine and Department of Biostatistics, University of Colorado, Denver, CO 80204, USA; Department of Medicine, University of California San Francisco, San Francisco, CA 94118, USA; CIBIO Research Center in Biodiversity and Genetic Resources, University of Porto, Vairo 4485-661, Portugal; Fundación Pública Galega de Medicina Xenómica-IDIS SERGAS, University of Santiago de Compostela, Santiago de Compostela, A Corua 15706, Spain; Section of Legal Medicine, Institute of Public Health, Catholic University of the Sacred Heart, Rome, RM 00168, Italy; Equipe des Sciences Anthropogénétiques et Biotechnologies, Faculté des Sciences, Universit Chouaib Doukkali, El Jadida, Doukkala Abda 24000, Morocco; Dipartimento di Biologia e Biotecnologie “Charles Darwin”, Sapienza Università di Roma, Rome, RM 00185, Italy; Laboratory of Human Genetics and Anthropology, Faculty of Pharmacy, University of Monastir, Monastir, Monastir 5000, Tunisia; Grupo de Medicina Xenómica-CIBERER-University de Santiago de Compostela, Santiago de Compostela, A Coruña 15782, Spain; Center of Excellence in Genomic Medicine, King Abdulaziz University, Jeddah, Jeddah 21589, KSA; Department of Bioengineering and Therapeutic Sciences, University of California San Francisco, San Francisco, CA 94118, USA; Department of Clinical Psychology and Psychobiology and Institute of Neurosciences, Universitat de Barcelona, Barcelona, Barcelona 08035, Spain; Institut de Recerca Sant Joan de Déu (IRSJD), Esplugues de Llobregat, Barcelona 08950, Spain; Section for Computational and RNA Biology, Department of Biology, University of Copenhagen, Copenhagen, Copenhagen 2100, Denmark

**Author notes:** These authors contributed equally to this work (Marc Via, Georgios Athanasiadis).

## Abstract

The Sephardim are a major Jewish ethnic division whose origins can be traced back to the Iberian Peninsula. We used genome-wide SNP data to investigate the degree of Sephardic admixture in seven populations from the Iberian Peninsula and surrounding regions in the aftermath of their religious persecution starting in the late 14^th^ century. To this end, we used Eastern Mediterranean (from South Italy, Greece and Israel) and North African (Tunisian and Moroccan) populations as proxies for the major ancestral components found in the target populations and carried out unlinked- and linked-marker analyses on the available genetic data. We report evidence of Sephardic ancestry in some of our Iberian samples, as well as in North Italy and Tunisia. We find the Sephardic admixture to be more recent relative to the Berber admixture following an out-of-Iberia geographic dispersal, suggesting Sephardic gene flow from Spain outwards. We also report some of the challenges in assigning Sephardic ancestry to potentially admixed individuals due to the lack of a clear genetic signature.

## Main text

The Sephardic Jews - or Sephardim - are a major Jewish ethnic division whose origins can be traced back to Spain and Portugal. There is documented presence of Jewish populations in the Iberian Peninsula and the Maghreb (North Africa, west of Egypt) since the Roman period^1^. The fate of the Sephardim waxed and waned over the centuries, from periods of tolerance and acceptance to eras of persecution, but the turning point occurred in 1391, when the Jews of Spain started to opt for conversion to Catholicism in increasing numbers as a response to violent attacks and murders. It is estimated that by early 15^th^ century, half of the approximately 400,000 Jews of Spain had already converted to Catholicism (becoming known as Conversos), while only about one-quarter remained as openly practicing Jews^2^. In 1492, the Catholic Monarchs of Spain issued the Alhambra Decree - an edict of expulsion whereby all Jews had to leave Spain by July of the same year.

The total number of Jews who left Spain has been a subject of controversy, although most recent estimates are between 50,000 and 80,000^3^. Part of the expelled Sephardic population settled in the Western Maghreb, whereas many others found shelter further east in the less hostile Ottoman Empire, which at that time included many Levantine and Balkan countries, as well as Turkey^4^. Sephardic communities were also established in other parts of Europe, such as France and Holland, as well as in Latin American countries^5^. Early on, many Spanish Jews went to Portugal, which at the time was more tolerant. However, in 1497, Portugal followed suit with its own Edict of Expulsion, forcing many Spanish and Portuguese Jews to leave Portugal or convert. It has actually been suggested that Portugal was less tolerant of their Jews to depart, and more were forced to stay in Portugal and live as Catholics^6^. Today, both Spain and Portugal offer citizenship to Jews of documented Sephardic descent.

By 1492, the number of Conversos remaining in Spain may have risen to 200,000 (or even 300,000) out of a total population of approximately 7.5 million and in early 16^th^ century, the proportion of the population of Sephardic origin is estimated to have been approximately 3–4%^3^. While soon after the Spanish and Portuguese Inquisitions Conversos were easily identified, their identity as former Jews was lost over time. Even though intermarriage between Conversos and Catholics was not commonplace at first, this changed over time, as the Jewish identity began to be lost. Therefore, one would expect to see evidence of Sephardic genetic ancestry among present day Iberians on a broad scale, and regionalization of the genetic signal is possible based on where the largest concentrations of Conversos had been living at the time of conversions.

Even though there have been many genetic studies on the origin of worldwide Jewish populations^7–11^, the genetic aftermath of the persecution of the Sephardim in the Iberian Peninsula is still poorly understood. A natural question to ask is the degree to which populations in the Iberian Peninsula show evidence of Sephardic genetic ancestry - and to what extent it may be regionalized. This question has been primarily addressed with uniparental markers^12,13^ - often with contrasting results. For instance, there is mtDNA-based evidence for Iberian admixture in Turkish Jewish communities^14^. Similarly, another mtDNA study showed that the Sephardim bore more resemblance to Spanish than to Jewish populations, but the opposite trend was observed for Y chromosomes^15,16^, a signal possibly reflecting the original Jewish colonization of Spain. On the contrary, the crypto-Jewish Xuetes from Majorca showed high frequency in R0, a mitochondrial lineage typically found in Middle Eastern populations - including the Jews^17^. Recent whole-genome studies have provided valuable insights into the genetic structure of the Iberian Peninsula^18^ (C.B., unpublished data). However, these studies lack at best an explicit reference to Sephardic genetic variation, limiting their scope primarily to North African genetic influences.

In light of the above, this study aims to provide a more detailed picture of the genetic influence of the Sephardim on a number of populations currently living in the Iberian Peninsula. Such influence would have probably been the result of Sephardic admixture into the majority Christian community through the forced conversions that started on a large scale in the late 14^th^ century, continuing through the time of the expulsion and beyond. To examine possible Sephardic admixture into other populations, we extended the geographic scope of our study to two Maghrebi populations from Morocco and Tunisia, as well as two European populations from South France and North Italy.

For this study, we used genotype data from 13 Mediterranean or near-Mediterranean populations of similar sample size (mean N ≈ 40; Table 1). More specifically, we typed 518 individuals (277 males and 241 females) on the Affymetrix 250K Sty array (Affymetrix, Santa Clara, CA, USA). The geographic origin of the samples is shown in Figure 1A. The Iraqi and Sephardic Jewish samples were collected in Israel and represent individuals with self-reported Iraqi Jewish or Turkish Sephardic grandparents, respectively. We included Iraqi Jews in our panel in order to account for Middle Eastern Jewish genomic ancestry. All participants provided written informed consent. Our study was conducted according to the Declaration of Helsinki II and was approved by the Bioethics Committees of the University of Barcelona, Spain and the University of California San Francisco, USA.

**Figure 1:**
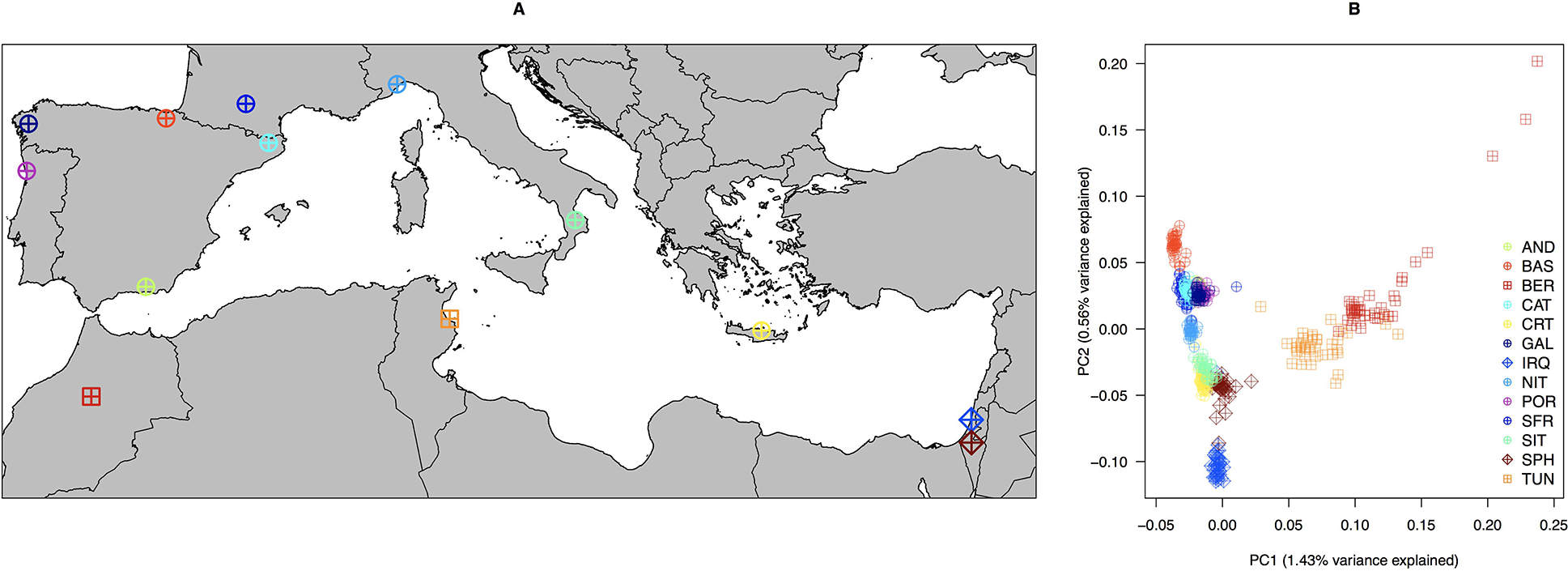
**(A)** Location of the 13 Mediterranean or near-Mediterranean populations used in this study (total N = 500). Circles plus represent European populations; squares plus represent African populations; and diamonds represent Israeli populations. **(B)** PCA based on 156,733 SNPs. Detailed information about geographic origin, abbreviations and sample sizes is shown in Table 1.

**Table 1:** Summary of the analyzed population samples Sample sizes are shown before and after quality control (QC), as well as after fineSTRUCTURE clustering.

| Country | Geographic region or ethnic group | Acronym | Initial N | N after QC | N after fineSTRUCTURE |
| --- | --- | --- | --- | --- | --- |
| Spain | Andalusians (Las Alpujarras) | AND | 40 | 40 | 40 |
| Spain | Basques (Gipuzkoa) | BAS | 37 | 37 | 37 |
| Spain | Catalans (Olot) | CAT | 45 | 41 | 39 |
| Spain | Galicians (Santiago de Compostela) | GAL | 34 | 34 | 34 |
| Portugal | Portuguese (Porto) | POR | 41 | 39 | 39 |
| France | South French (Toulouse) | SFR | 46 | 46 | 44 |
| Italy | North Italians (Liguria) | NIT | 31 | 29 | 28 |
| Italy | South Italians (Calabria, Puglia, Campania & Basilicata) | SIT | 35 | 35 | 35 |
| Greece | Cretans (Crete) | CRT | 44 | 44 | 44 |
| Morocco | Berbers (Khenifra) | BER | 40 | 39 | 38 |
| Tunisia | Tunisians (Monastir) | TUN | 41 | 39 | 36 |
| Israel | Iraqi Jews | IRQ | 42 | 38 | 37 |
| Israel | Sephardic Jews (Turkish origin) | SPH | 42 | 39 | 39 |
|  |  | **Total** | **518** | **500** | **490** |

Standard quality control was performed with PLINK v1.9^19^ and included the removal of duplicates, close relatives, extreme outliers, as well as individuals with >5% genotype missingness. We subsequently removed variants with >5% genotype missingness and those that did not fit per-population Hardy-Weinberg proportions after Bonferroni correction. After quality control, a total of 500 samples and 214,338 autosomal single nucleotide polymorphisms (SNPs) were available for downstream analyses.

We performed principal component analysis (PCA) on all 13 populations with PLINK using a set of 156,733 SNPs obtained after linkage disequilibrium (LD) pruning (*r*^*2*^ threshold = 0.8; window size = 50; step size = 5). The first PC separates North Africans (Berbers and Tunisians) from the rest of the populations, whereas the second PC roughly corresponds to the distribution of the remaining populations along an east-west axis with the Iraqi Jews and the Basques occupying the two extremes (Figure 1B). The PCA plot revealed that the Sephardic Jews are genetically most similar to Eastern Mediterranean populations, such as Cretans and South Italians, and to a lesser degree to the Iraqi Jews, in agreement with previous observations^7,8^. The observed genetic affinities remained the same after first removing the North Africans (Figure S1), and then the Eastern Mediterraneans and Jewish populations (Figure S2) from the PCA.

We also ran an unsupervised ADMIXTURE analysis^20^ using the same SNP set as in PCA. ADMIXTURE corroborated our PCA findings regarding the genetic relationship between the Jewish populations and the rest of the samples (Figure 2). For K = 3 (i.e. the model with the smallest cross-validation error = 0.457), Iraqi Jews, Basques and Berbers appear as the most differentiated and homogeneous clusters (for comparison see Figure 1B). For K ≥ 4, the Iraqi Jews persist as a largely homogeneous cluster, whereas for K = 7 in particular, the Sephardic Jews acquire their own cluster (in brown), which we can then trace in other populations. This putatively Sephardic component was notably present in Tunisia (27.23%), but also in the Iberian Peninsula with the exception of the Basque Country (Galicia: 4.25%; Portugal: 2.39%; Andalusia: 2.36%; Catalonia: 1.32%), South France (1.56%), North Italy (1.70%) and South Italy (7.50%).

**Figure 2:**
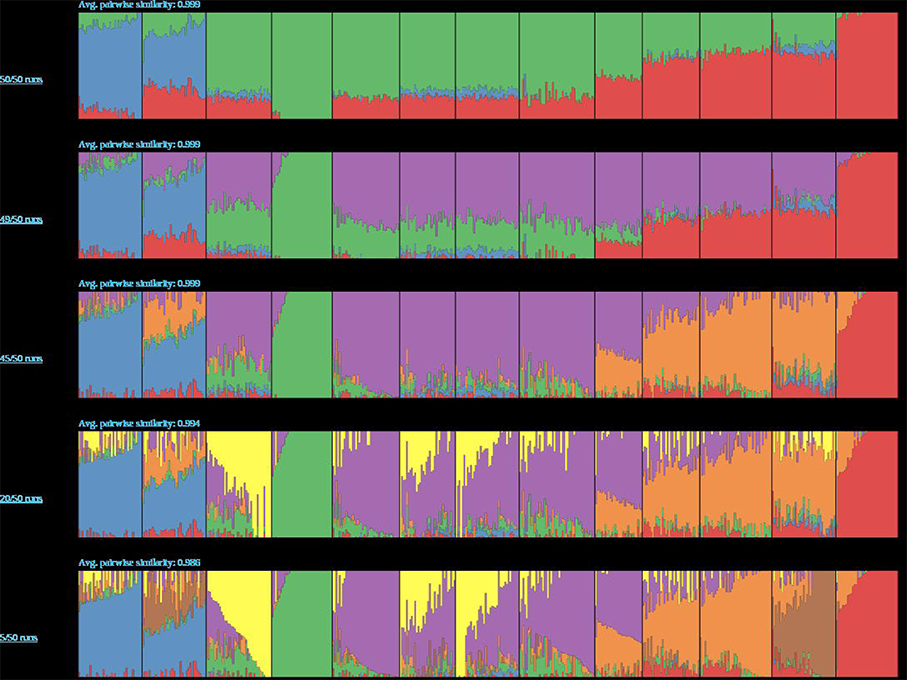
Ancestral component analysis of the 13 Mediterranean or near-Mediterranean studied populations assuming K = {3, 4, 5, 6, 7} admixing populations. ADMIXTURE was run 50 times for each K and results were plotted with PONG^31^. Bar plots show the consensus per-individual membership to each of the specified ancestral components. Population labels are described in Figure 1.

We then used CHROMOPAINTER, fineSTRUCTURE and GLOBETROTTER^21,22^ in their default settings to achieve a more detailed picture of population structure in our samples. Because our SNP chip was of an older generation, we used liftOver from the University of California Santa Cruz Genome Browser to update SNP positions from the NCBI36/hg18 (March 2006) to the GRCh37/hg19 build (February 2009). To increase the total number of available SNPs necessary for these analyses, we carried out genotype imputation using the Michigan Imputation Server^23^ (phasing software: Eagle v2.3^24^; imputation software: Minimac v3^23^; reference panel: hrc.r1.1.2016; QC’s reference population: mixed population). After filtering for info ≥ 0.995 and removing monomorphic loci and singletons, there were 527,379 phased autosomal SNPs for the painting analyses.

We first carried out a painting in which each of the 500 samples was allowed to copy DNA segments from all of the remaining samples. CHROMOPAINTER returns similarity matrices of shared haplotype counts and shared haplotype lengths. We used the count matrix together with fineSTRUCTURE to hierarchically cluster the 500 individuals into a Bayesian consensus phylogenetic tree. fineSTRUCTURE returned a tree of nine largely homogeneous clusters (Figure S3), which resemble to some extent the results from PCA (Figure 1B) and ADMIXTURE (Figure 2). As before, the most differentiated clusters were the ones from North Africa (Tunisians and Berbers), whereas the next node divided the remaining populations into two clusters: Southwest vs. Southeast Europe (including the two Jewish populations). The clustering is consistent with geography with the exception that North Italy was clustered with the Southwestern populations rather than Southeastern. South Italy and Crete clustered together, and the Sephardic Jews appeared more closely related to these two populations than to the Iraqi Jews. To add more resolution, we repeated the painting using only the Iberian (without the Basques) and French populations, in which as above each individual was allowed to copy DNA segments from all other individuals. In this case, fineSTRUCTURE returned a tree with four overly homogeneous clusters, in which the Galician and Portuguese were the only samples that did not split according to their labels (and were therefore treated as one group in the subsequent analyses), while Catalonia, South France and Andalusia formed virtually homogeneous groups (Figure S4).

We used the information from the two fineSTRUCTURE analyses to select recipient and donor groups and carried out two additional paintings in which (i) recipients were allowed to copy DNA segments only from donors (i.e. donor-to-recipient painting) and (ii) each donor was allowed to copy DNA segments only from other donors (i.e. donor-to-donor painting). For this particular analysis, we used CHROMOPAINTER’s length matrix together with GLOBETROTTER to estimate South European, Sephardic and North African (i.e. donors) admixture proportions in the Iberian, French, North Italian (i.e. recipients) clusters.

Figure 3A shows the main Mediterranean donor group contributions to six Southwestern European clusters. Among these, the highest Sephardic admixture appeared in Andalusia (12.3%; 95%CI: 11.1-13.5%), followed by Galicia and Portugal (11.3%, 95%CI: 10.6-12.1%), while it was 5.9% in North Italy (95%CI: 5-6.8%). The Sephardic component was notably absent in the Basque Country and South France, as well as in Catalonia (point estimate of ~0.3%, not statistically different from zero). These results reflect the regionalization of Sephardic admixture on the Iberian map and surrounding regions. Our Sephardic admixture estimates in the Iberian Peninsula were overall more conservative than those reported elsewhere for Y chromosome data^25^ (their mean value of Sephardic ancestry: 19.8% vs. ours: 6%). The Iraqi Jews were the only donor group that did not contribute to any of the recipient clusters, while the Berber component was present in all of the Iberian populations (with the exception of Catalonia) and South France, matching historical knowledge about the Moorish presence in the Iberian Peninsula during the Middle Ages^26^, as well as results from a recent study in the same region (C.B., unpublished data). In addition, the higher Sephardic ancestry in Galicia and Portugal compared to the rest of the peninsula could be reflecting the ban on the departure of Jews from the kingdom of Portugal combined with greater pressure to convert to Catholicism (thus corroborating historical hypotheses) and/or Sephardic gene flow from Spain to Portugal^27,28^.

**Figure 3:**
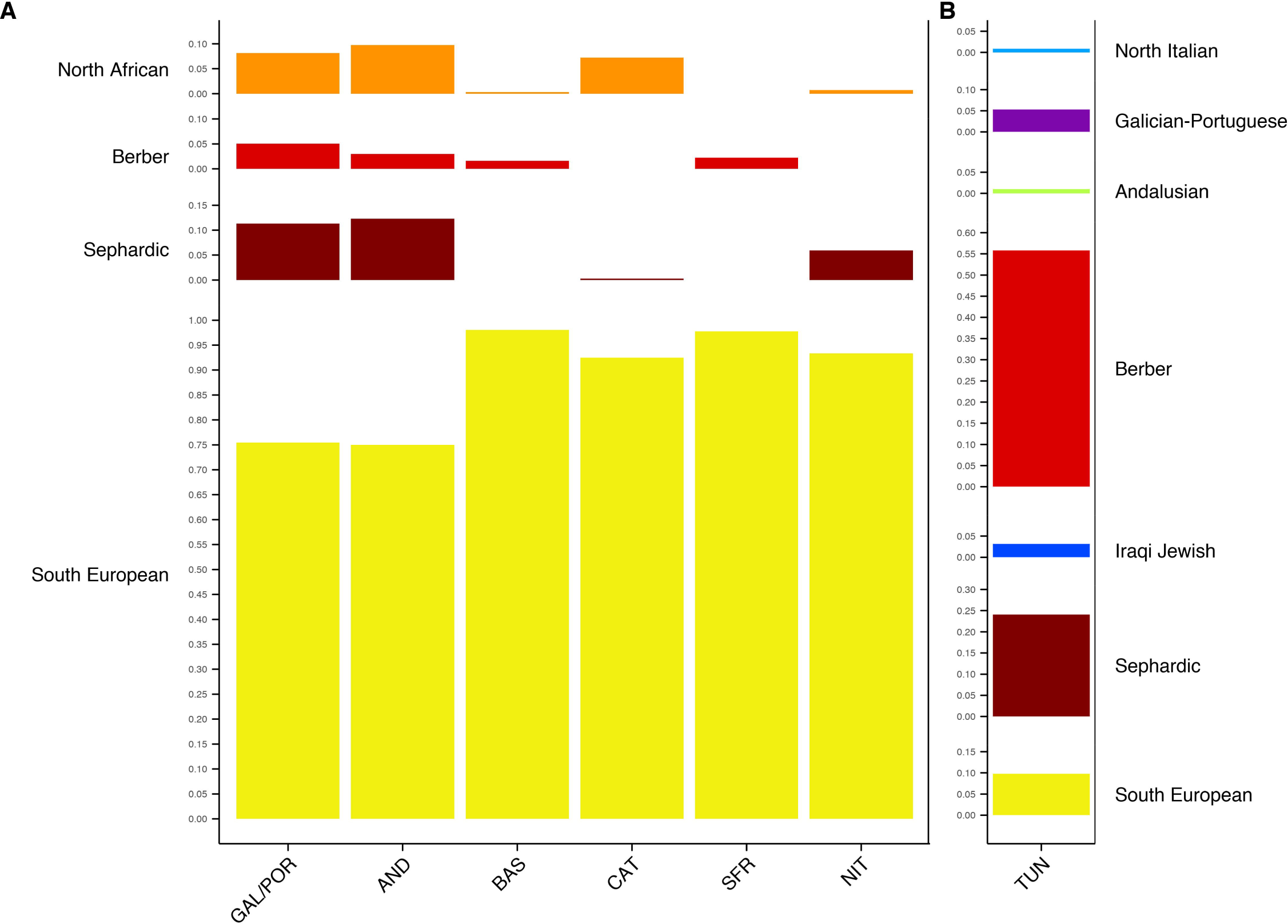
**(A)** GLOBETROTTER admixture profiles for six populations from the Iberian Peninsula, France and Italy. Of the five admixture components used in the model (South European, Berber, North African, Sephardic and Iraqi Jewish), only the first four contributed substantially to the recipient populations. The South European component included samples from South Italy and Crete. **(B)** GLOBETROTTER admixture profile for the Tunisian population using seven admixture components (South European, Sephardic, Iraqi Jewish, Berber, Andalusian, Galician-Portuguese and North Italian). Population legends are described in Figure 1.

Historical records show that many Sephardic Jews sought shelter in the Maghreb after they were expelled from Spain and Portugal^1^. In light of this, we performed an additional CHROMOPAINTER and GLOBETROTTER analysis, this time using Tunisians as the recipient population, and all of the remaining clusters as donors, in order to estimate the Sephardic influence in North Africa (Figure 3B). The analysis showed that the main ancestry contributors for Tunisia were the Berbers (55.8%; 95%CI: 55.3-56.3%) followed by the Sephardic Jews (24.1%; 95%CI: 23.2- 25%), and to a lesser extent by the remaining donor clusters. Even though the Sephardim were present in Tunisia, it is unlikely that their genetic contribution is as high as reported here by either ADMIXTURE or GLOBETROTTER. The rate of conversion of Sephardic Jews into the local Muslim population was low and intermarriage between Jews and Muslims has been limited throughout their history^29^. Rather, we attribute this high percentage to lacking an appropriate reference population that reflects better the historical background of Tunisia (e.g. a proxy population for the Phoenicians). In Antiquity, the Phoenicians not only had a major capital in Tunisia (Carthage), but also significant outposts in the Iberian Peninsula. As a Levantine population, the Phoenicians were probably genetically very similar to the ancestral Jewish populations of the Mediterranean. Bearing this in mind, we note that our Sephardic estimates for the Iberian Peninsula are potentially also liable to the same issue and should therefore be interpreted as an upper bound.

Finally, in order to provide temporal context for our observations in Southwestern Europe, we inferred local ancestry on phased chromosomes using RFMix^30^. RFMix takes into account LD among markers and identifies ancestry tracts originating in each of the chosen admixing populations. For this analysis, target populations included only samples with considerable Sephardic ancestry, i.e. Andalusia, Galicia-Portugal and North Italy. We inferred local ancestry patterns on target populations as the result of a three-way admixture between Sephardim, Berbers and Iberians. Sephardic genetic variation was represented by the Sephardic Jews (N = 39); Berber variation by the Moroccan Berbers (N = 38); and Iberian variation by 10 samples from Andalusia and Portugal that showed high membership (≥99%) to the yellow cluster for K = 7 in Figure 2. We ran RFMix three times, assuming time of admixture g = {10, 25, 40} generations ago. The patterns of tract length distribution were qualitatively similar across all three choices of g (differing only in scale), so we chose to focus the rest of our report on g = 25 (~625 years before present).

Figure 4A shows the distribution of migrant tract lengths for the pooled Iberian populations (Andalusia and Galicia/Portugal), whereas Figures S5-7 show the same distribution for each population and each time of admixture g separately. In all cases, Sephardic tracts were predominant over Berber tracts, both in frequency and in length, implying more recent admixture with the Sephardim than with the Berbers.

**Figure 4:**
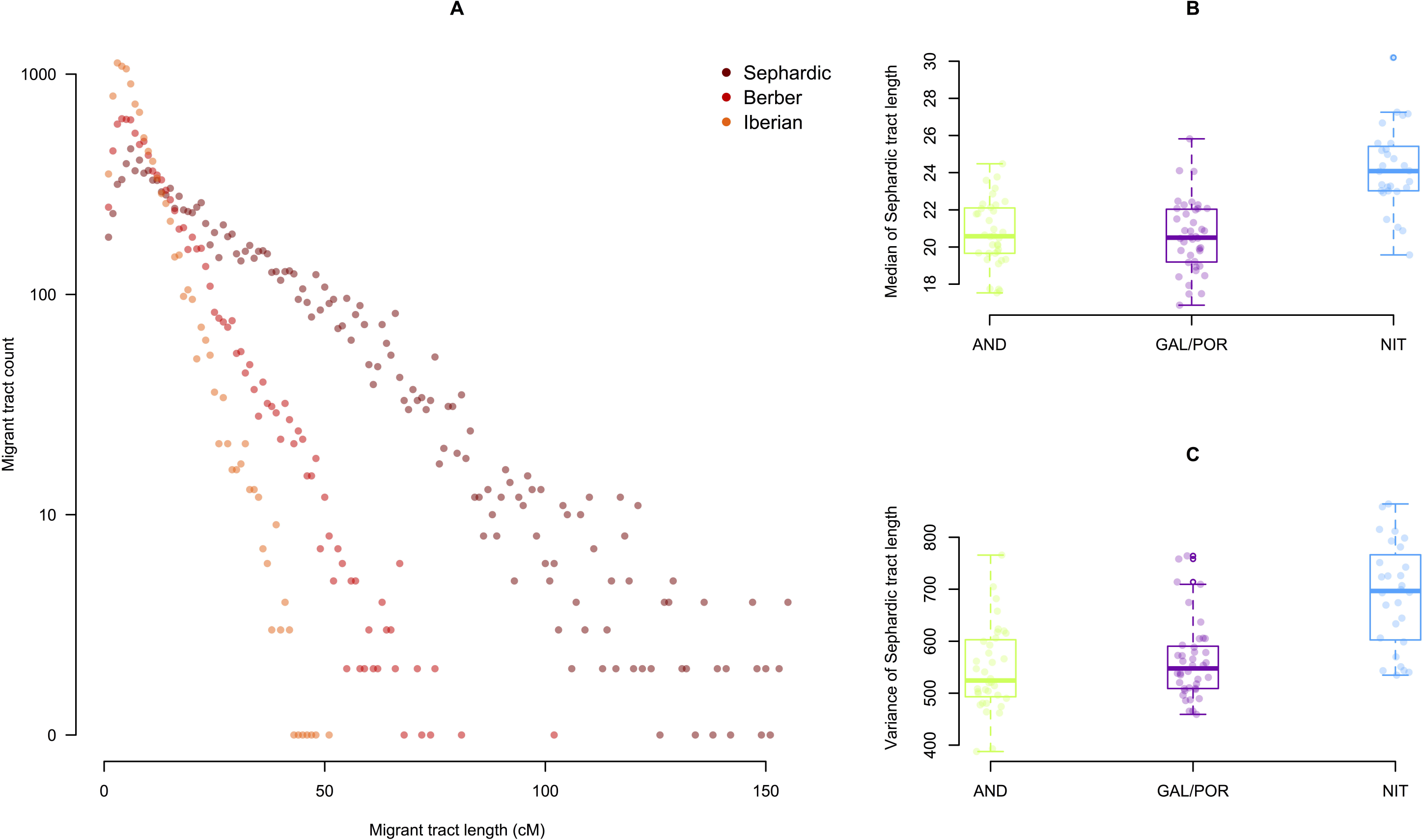
**(A)** Distribution of Sephardic, Berber and Iberian migrant tracts (1-cM bins) in a pooled sample made up of Andalusians and Galician-Portuguese assuming time of admixture g = 25 generations ago. Migrant tract length was calculated by subtracting initial from final genomic position (in cM) of each tract as reported by RFMix. Tract length calculations were restricted within chromosome arms, thus omitting centromeres. Berber tracts were notably shorter than Sephardic tracts, suggesting that Berber admixture is older than Sephardic admixture in the Iberian Peninsula. **(B)** Jitter and box plot of (i) median Sephardic tract length and (ii) variance of Sephardic tract length for Andalusians, Galician-Portuguese and North Italians. Sephardic migrant tracts look younger (i.e. with larger median and variance) outside the Iberian Peninsula. Population legends are described in Table 1.

We then checked whether Sephardic tract length for each of the three populations under study correlated with geography. Figures 4B-C show the relationship (i) between geographic location and median Sephardic tract length, and (ii) between geographic location and variance in Sephardic tract length for the three target populations, whereas Figure S8 shows the same information for g = {10, 40}. Even though the effect is subtle, we detected a geographic pattern, with ancestry tracts looking younger (higher median length and variance) outside Iberia and older (lower median length and variance) inside Iberia. Combined with the relative tract length distribution (Figure 4A), we interpret these observations as suggestive of recent Sephardic gene flow from the Iberian Peninsula towards the East of the Mediterranean. In addition, tract lengths in the two Iberian populations (i.e. Andalusia and Galicia-Portugal) followed very similar distributions (Figures 4B-C), probably reflecting near-contemporary admixing events.

Recent whole-genome studies on the structure of worldwide Jewish groups showed that current Sephardic populations are more related to other Jewish and Middle Eastern groups than to non-Jewish Spaniards and Northwest Africans^7–9^. Here, we detected a Sephardic component in the Iberian Peninsula, as well as North Italy and Tunisia. A possible reason for this discrepancy could be the methods used; we observed that PCA did not detect the Sephardic genetic influence in the Iberian Peninsula (Figures 1B and 2), yet haplotype-based methods such as CHROMOPAINTER and RFMix, were more powerful at picking up signatures of Sephardic admixture (Figures 3 and 4). We therefore recommend haplotype-based methods for the discovery of Sephardic admixture signals in follow-up studies.

An interesting question regarding the era of the Unification of Spain is whether admixture between the Christian, Muslim and Jewish groups was mainly ancient or promoted by the persecution and consecutive conversion. The relative frequency of migrant tracts suggests that Sephardic admixture is notably more recent than Muslim admixture. Given that the presence of Sephardic populations in the Peninsula predates the Arab invasion, this can be interpreted as Sephardic admixture occurring in more recent times, perhaps as intermarriage under the newly acquired Catholic label. An additional hint that corroborates this idea comes from the distribution of Sephardic tracts along Southwestern Europe, as Sephardic gene flow seems to follow an out-of-Iberia pattern, being more recent outside the Iberian Peninsula, although more samples from their potential route to the East are warranted to draw clearer conclusions.

The high percentage of Sephardic ancestry in Tunisia (and probably also Northern Italy, even though the numbers there are quite smaller), as reported both by ADMIXTURE and GLOBETROTTER, does not match historical expectations. We believe that this could be due to the lack of a more appropriate Levantine donor population for the painting of Tunisian chromosomes. This observation hints at the lack of sufficient specificity in the Sephardic genetic signature, probably reflecting a less tumultuous demographic history for the Sephardim compared to e.g. the Ashkenazim. We therefore interpret our admixture estimates as an upper bound.

In this study, we reassessed Sephardic ancestry in the genetic pool of the Iberian Peninsula and parts of Northwest Africa. To do so, we used a sample of Israelis with all four of their grandparents of Turkish Sephardic descent as a proxy for a Sephardic-descended population. In our analyses, Sephardic ancestry was present in many but not all of the studied samples from the Western Mediterranean, possibly reflecting the regionalization of Sephardic admixture and the historical particularities of each region. The observed geographic patterns serve as a starting point to test more specific hypotheses about the course of events around the time of the persecution of the Sephardim in the Iberian Peninsula. Future research should include a broader geographic scope encompassing not only the Mediterranean but also other parts of the globe (J.C., unpublished data) and, ideally, denser SNP arrays.

## Acknowledgements

We would like to thank all participants that provided samples for the study. Genotyping services were provided by the Spanish “Centro Nacional de Genotipado” (CEGEN-ISCIII)” and we also thank the CEGEN coordination team for their support. E.G.B. is supported by the Sandler Family Foundation, the American Asthma Foundation, the RWJF Amos Medical Faculty Development Program, the Harry Wm. and Diana V. Hind Distinguished Professor in Pharmaceutical Sciences II, the National Heart, Lung, and Blood Institute (R01HL117004, R01HL128439, R01HL135156, X01HL134589), the National Institute of Environmental Health Sciences (R01ES015794, R21ES24844), the National Institute on Minority Health and Health Disparities (P60MD006902, R01MD010443, RL5GM118984) and the Tobacco-Related Disease Research Program (24RT-0025).

## Figure titles and legends

**Figure S1:** PCA based on 156,733 SNPs for the South European and Jewish populations (i.e. without the North Africans).

**Figure S2:** PCA based on 156,733 SNPs for the Iberian Peninsula, South France and North Italy.

**Figure S3:** fineSTRUCTURE grouping of 500 Mediterranean or near-Mediterranean samples roughly corresponding to well-defined geographic locations.

**Figure S4:** fineSTRUCTURE grouping of 198 Iberian (without the Basques) and South French samples roughly corresponding to well-defined geographic locations. Only Galicia and Portugal did not form clear clusters according to their labels.

**Figure S5:** Distribution of Sephardic, Berber and Iberian migrant tracts (1 cM bins) in Andalusians, Galician-Portuguese and North Italians assuming time of admixture g = 10 generations ago.

**Figure S6:** Distribution of Sephardic, Berber and Iberian migrant tracts (1 cM bins) in Andalusians, Galician-Portuguese and North Italians assuming time of admixture g = 25 generations ago.

**Figure S7:** Distribution of Sephardic, Berber and Iberian migrant tracts (1 cM bins) in Andalusians, Galician-Portuguese and North Italians assuming time of admixture g = 40 generations ago.

**Figure S8:** Jitter and box plot of (i) median Sephardic tract length and (ii) variance of Sephardic tract length for Andalusians, Galician-Portuguese and North Italians, assuming time of admixture g = {10, 40} generations ago.

